# Fast 3D imaging of giant unilamellar vesicles using reflected light-sheet microscopy with single molecule sensitivity

**DOI:** 10.1101/2020.06.26.174102

**Authors:** Sven A. Szilagyi, Moritz Burmeister, Q. Tyrell Davis, Gero L. Hermsdorf, Suman De, Erik Schäffer, Anita Jannasch

**Author notes:** contributed equally (Sven A. Szilagyi, Moritz Burmeister). http://www.zmbp.uni-tuebingen.de/res/nano.

## Abstract

Observation of highly dynamic processes inside living cells at the single molecule level is key for a quantitative understanding of biological systems. However, imaging of single molecules in living cells usually is limited by the spatial and temporal resolution, photobleaching and the signal-to-background ratio. To overcome these limitations, light-sheet microscopes with thin selective plane illumination have recently been developed. For example, a reflected light-sheet design combines the illumination by a thin light-sheet with a high numerical aperture objective for single-molecule detection. Here, we developed a reflected light-sheet microscope with active optics for fast, high contrast, two-color acquisition of *z*-stacks. We demonstrate fast volume scanning by imaging a two-color giant unilamellar vesicle (GUV) hemisphere. In addition, the high signal-to-noise ratio enabled the imaging and tracking of single lipids in the cap of a GUV. In the long term, the enhanced reflected scanning light sheet microscope enables fast 3D scanning of artificial membrane systems and cells with single-molecule sensitivity and thereby will provide quantitative and molecular insight into the operation of cells.

## 1. Introduction

The ability to image and track single molecules in living cells in real time and three-dimensions (3D) represents one of the big challenges in microscopy. A powerful approach to improve the precision of single molecule localization, is to maximize the collection of emitted photons and reduce the background signal from out-of-focus fluorophores. Collection efficiency scales quadratically with the numerical aperture (NA) of the imaging objective. Thus, the NA should ideally be as large as possible. Total internal reflection fluorescence microscopy (TIRFM), widely used for single molecule studies, illuminates only a thin plane near the glass-sample interface. This reduction in illumination volume results in a high signal-to-noise ratio (SNR) of single fluorophores. However, due to the intrinsically restricted geometry of illumination, TIRFM is not suitable for 3D imaging of whole cells. To achieve a high SNR for 3D imaging, different designs of light-sheet microscopes have been developed [1, 2]. Light-sheet illumination reduces the out-of-focus background, resulting in a better SNR. Most importantly, the intrinsic optical sectioning reduces phototoxicity, photobleaching and enhances imaging speed. Therefore, fast 2D and 3D imaging can be realized making light-sheet microscopes a powerful tool for imaging live cells. Traditional light-sheet microscopes (*e.g.* SPIM) were designed for measuring the real-time dynamics of relatively large samples, such as embryos [3, 4]. In these microscopes, the illumination objective is placed perpendicular to the detection objective. However, due to spatial constraints, high NA objectives with an inherent small working distance cannot be used which limit this design. Sub-cellular high-resolution and high-contrast imaging requires a thin light-sheet illumination and the usage of high NA detection objectives. Various optical designs have been made to combine single-molecule super-resolution with different approaches of light-sheet illumination [5, 6]. Digitally scanned laser light-sheets, (*e.g.* Bessel beam light-sheet) improve the uniformity of the light-sheet profile and increase the axial resolution [7–9]. The tilted light-sheet microscopy with 3D point spread functions combines a tilted light-sheet illumination strategy with long axial range point spread functions. This approach allows one color 3D super-localization of single molecules [10]. Single objective light-sheet microscopes (*e.g.* HILO, soSPIM,) use high NA objectives for both illumination and detection. The oblique illumination of the HILO microscope limits the illumination area, which decreases proportionally to the light-sheet thickness [11]. Other single objective light-sheet microscopes use reflective surfaces or prisms, requiring custom-made sample chambers or have limited multi-color imaging capabilities [12–15]. The reflected light-sheet microscope combines thin light-sheet illumination with a parallel arrangement of illumination and detection objectives [16–18]. The light-sheet is reflected by a small cantilever mirror into a horizontal plane close to the sample surface. This design allows the use of a high NA objective for horizontal sectioning of samples with a light-sheet thickness of ≈ 1 μm, as well as a high NA objective for detection of fluorescence.

We adapted the principle of the reflected light-sheet and further developed the design for fast 3D imaging with single molecule sensitivity. Fast 3D scanning of the sample is realized by implementation of active optics allowing the light-sheet to be automatically moved through the specimen without moving the sample stage [19, 20]. The active optics enable fast 3D scanning of the thin light-sheet over a volume of 35 × 10 × 15 μm^3^ (*x, y, z*) in less than 500 ms. For simultaneous two color imaging, we implemented two lasers (488 nm and 561 nm) and dual-view imaging optics. In addition, a 405 nm light-sheet was co-aligned for (re-)activation of fluorophores. Thus, in principle, photoactivation experiments with single-molecule 3D super-resolution imaging can be performed in thick biological samples, such as whole cells [16, 21]. Here, we demonstrate the imaging and detection of single fluorophores diffusing in a giant unilamellar vesicle (GUV) and show a fast two-color, 3D scan of a GUV.

## 2. Material and methods

### 2.1. Design of the reflected 3D scanning light-sheet microscope

The custom-build reflected light-sheet microscope is built on an optical table with a custom designed damping system [22]. To avoid drift and diffusive light from any light source other than the illumination source, all adjustments during an experiment are controlled via motorized components from outside the dark inner microscope room. In addition, to visualize the sample, a bright field and interference reflection microscope (IRM, [23, 24]) are included (Fig. 1).

**Fig. 1.**
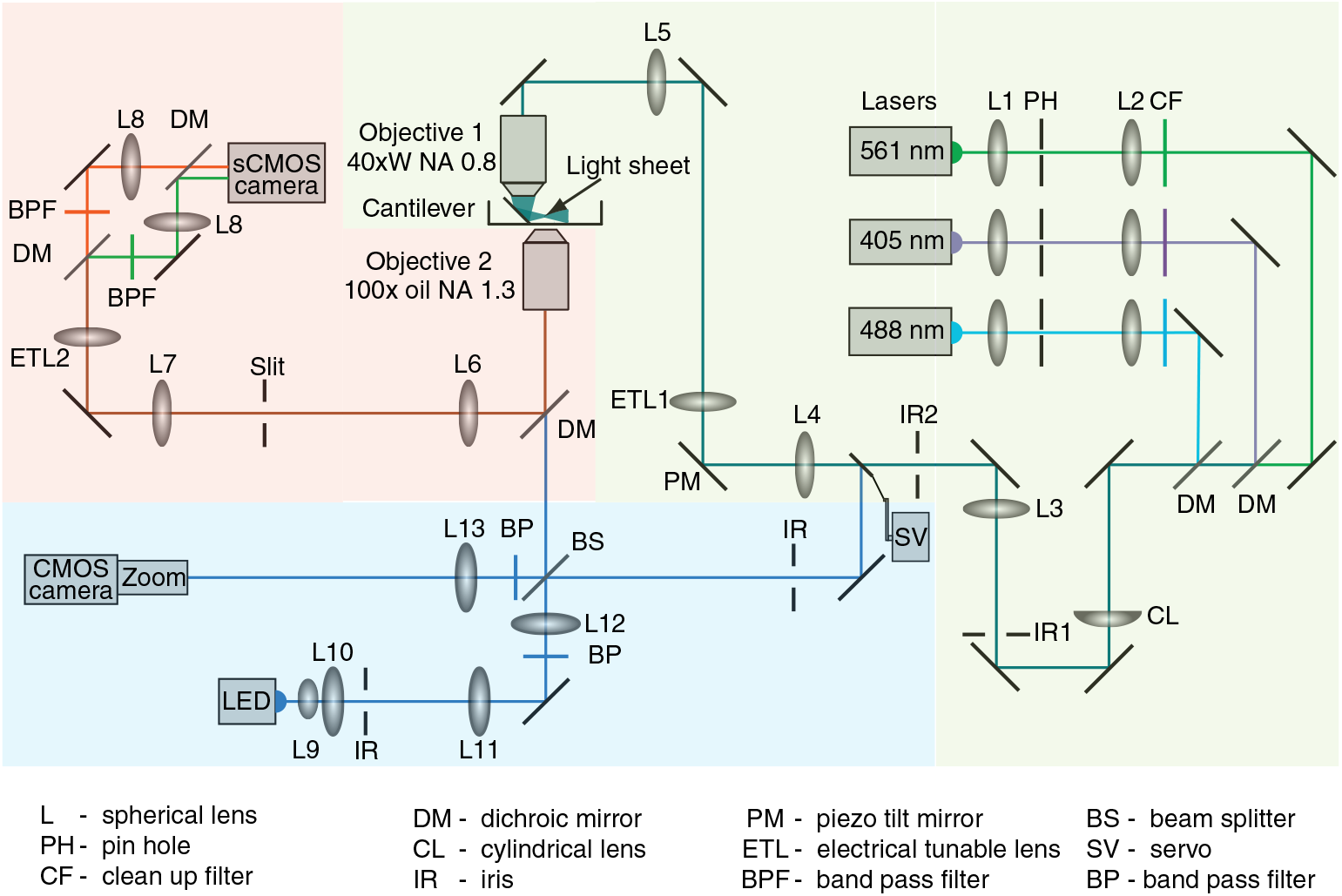
Schematic of the reflected 3D scanning light-sheet microscope design. Shown are the fluorescence excitation and generation of the thin light sheet (light green box), the fluorescence detection path (light red box), and the interference reflection microscopy and bright field illumination path (light blue box). The optical path is drawn to scale. Focal length of lenses L1–L13 in mm are: 25, 100, 100, 100, 200, 160, 80, 150, 20, 40, 80, 300, and 200, respectively.

#### 2.1.1. Fluorescence excitation and generation of the thin light sheet

For fluorescence excitation and photoactivation, three diode lasers (405 nm: LuxX 405-120, Omicron-Laserage; 488 nm: LuxX 488-100, Omicron-Laserage; 561 nm: OBIS 561LS-100, Coherent) are collimated and aligned via telescopes (Lens L1, L2, Qioptiq Photonics, Germany), pinholes (PH,10 μm, Qioptiq Photonics, Germany), and dichroic mirrors (AHF Analysentechnik, Germany). Note that the laser power is significantly reduced by the pinholes. The laser beams are further expanded and focused by a cylindrical lens (CL, *f* = 150 mm, Thorlabs) yielding a line profile in the back focal plane of the high NA, water dipping illumination objective (Objective 1, CFI Apo NIR 40 × W, 0.8 NA, Nikon, Japan). The cylindrical lens is placed on a motorized rotation mount (SR-7012-S, SmarAct, Germany) allowing different orientations of the light-sheet. To control the width and the thickness of the light-sheet, two irises (IR1, IR2) are employed. After passing the illumination objective, the thin light-sheet is reflected by a tipless gold-coated atomic-force-microscopy cantilever (HYDRA2R-100NG-TL, AppNano, USA). The backside of this cantilever acts a mirror reflecting the light-sheet by 90◦ parallel to the imaging plane. The cantilever is attached to piezo positioners (SLC-1720, SR-7012, SmarAct, Germany) that enable 3D movement as well as rotation around the illumination objective. In this manner, an object of interest can be illuminated from a desired orientation. In addition, a piezo tilt mirror (PM, TT2.5, Piezoconcept, France) is placed in the conjugated plane of the back focal plane of the illumination objective (Objective 1) enabling fast movement of the reflected light-sheet in the *z*-axis of the sample plane. An electrically tunable lens (ETL1, EL-10-30-C-VIS-LD, Optotune, Switzerland) is used to adjust the focus position of the light-sheet relative to the cantilever mirror and sample of interest. The piezo motors and ETLs are controlled via LabView (National Instruments, Austin, Texas, USA).

#### 2.1.2. Fluorescence detection

The fluorescence emission is collected by a high NA oil immersion objective (Objective 2, CFI S Fluor 100 × Oil, 1.3 NA, Nikon, Japan) and imaged onto a sCMOS camera (ORCA-Flash4.0 V2, Hamamatsu, Japan). With the overall magnification, the image pixel size corresponded to 26 nm. With a second ETL (ETL2, EL-16-40-TC-VIS-5D, Optotune, Switzerland), the *z* position of the image plane inside the sample above the detection objective can be adjusted. Note that both ETLs are mounted with their optical axis parallel to the gravitational acceleration vector. Simultaneous image acquisition of two colors is realized by image-splitting optics (dichroic mirror DM, band pass filter BPF, Lens L8) placed in front of the camera [23]. Through individual placement of Lens 8, the color splitter also allows adjustment to color-specific focal planes. The lasers and the camera are triggered and controlled via μManager [25].

#### 2.1.3. Interference reflection and bright field microscope

For the LED-based IRM [23], an image of the LED (450 nm Royal-Blue LUXEON Rebel LED, Lumileds, Germany) is magnified by two telescopes (Lens L9-12) and projected into the back focal plane of the detection objective (Objective 2). The detection and illumination light paths are separated by a 50/50 beam splitter plate (BS, F21-000, AHF Analysentechnik, Germany). A tube lens (Lens L13) and a zoom (S5LPJ7073, Sill Optics, Germany) magnify the sample image about 70× and project it onto a CMOS camera (Lt225, Lumenera, USA). Additionally, the LED is used for bright field illumination by coupling the light via the BS and a movable mirror that is mounted on a servo and can be switched in and out of the illumination path of the light sheet. Thus, while IRM and light-sheet based imaging can be done simultaneously, brightfield microscopy is only possible without light-sheet illumination. In the setup, brightfield microscopy is useful for initial alignment of the cantilever.

### 2.2. Preparation of GUVs and supported lipid bilayers

The GUVs were prepared using established protocols for electroformation [26, 27]. Briefly, 2 μl of a lipid mixture (99.95 mol% DOPC, 0.044 mol% DiO (3,3’-dioctadecyloxacarbocyanine perchlorate), 3 × 10^−7^ mol% rhodamine-PE) in chloroform were pipetted on each of the two platinum wires in the lid of a custom-made chamber made out of polytetrafluoroethylene. The lipid mixture was dried under vacuum for 10 min and the chamber filled with 350 μl of a 300 mM sucrose solution. The lid was screwed on the chamber and the platinum contacts were connected to a function generator (H-Tronic, FG 250 D). For electroformation of GUVs, we used a frequency of 10 Hz with a 2 V amplitude over a time of 60 min. Release of GUVs from the electrodes was achieved with a frequency of 2 Hz and 2 V amplitude for 30 min. Before experiments, the glass surface of a glass bottom petri dish (ibidi, μ-Dish 35 mm) was blocked with 50 μl BSA (18 mg/ml). Afterwards, the petri dish was filled with 1 ml of a 310 mM glucose solution supplemented with an oxygen-scavenger system (20 μg/ml glucose oxidase, 8 μg/ml catalase). Finally, the sucrose GUV solution was added and allowed to settle to the glass surface for approximately 30 min. The supported lipid bilayers were prepared using vesicle deposition [28, 29]. The lipid mixture and dye ratio were the same as for the GUV preparation.

### 2.3. Data acquisition and analysis

For light-sheet characterization, a mixture of 1 μg/ml fluorescein and 1 μg/ml tetramethylrhodamine in 2 ml Milli-Q water was used. Images were acquired using 2 × 2 binning of the sCMOS camera resulting in an effective pixel size of 52 nm. The exposure time (10–100 ms) and the laser power (1–100 mW) were varied to find an optimal balance between signal intensity and temporal resolution. For all images, background images were taken and subtracted. The images were analyzed using the image processing package Fiji [30]. Experiments were carried out at room temperature (25 °C).

For tracking of single fluorophores, the Trackpy (v0.4.1) package for Python was employed [31]. The Gaussian spot localization in the GUV images was carried out with a spot diameter of 9 pixels and minimal integrated brightness of 150 counts. The detected features in each image were then linked to subsequent ones in the image stack with a maximum travel distance of 10 pixels per frame and at most five skipped frames. The resulting trajectories were filtered to have a length of at least 30 frames or more.

The 3D model of the GUV was reconstructed via the 3D Viewer plugin in Fiji with a threshold of 50 and a resampling factor of 4 for the surface plot and a resampling factor of 1 for the volume plot [30].

## 3. Results and Discussion

### 3.1. Characterization of the light-sheet

The light-sheet setup enables fast 3D imaging of GUVs and single cells using a thin light-sheet reflected by a gold-coated atomic-force-microscopy cantilever placed next to the sample of interest. The custom-built setup is schematically illustrated in Fig. 1 and described in detail in the Methods. To quantify the optical properties of the light-sheet, we either imaged the lasers directly or excited a fluorophore solution using the rotatable light sheet with and without cantilever (Fig. 2, see Methods). By projecting the light-sheet focus into the image plane of the high NA detection objective without the cantilever, fluorophores, and emission filters, we could directly image the cross section of the lasers that form the light sheet and confirm that all three lasers were coaligned (Fig. 2a). Using Iris 2 (Fig. 1), we adjusted the width of the light sheet to match the width of the cantilever (35 μm). By inserting the cantilever mirror, the light sheet is reflected into a plane coinciding with the imaging plane. Using a fluorescent solution excited by the 488 nm and 561 nm lasers (Fig. 2b), we could verify both the width and even illumination of the excited region. To determined the thickness of the light sheet during normal imaging, we rotated the cylindrical lens by 90 degrees and imaged the excited cross section normal to the plane of the light sheet (Fig. 2c). The intensity distribution of the most narrow light-sheet cross section was then analyzed by fitting Gaussian functions 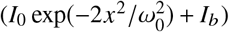 through the cross sections, where ω_0_ is the thickness of the light-sheet and *I_b_* is the intensity of the background signal (Fig. 2d). The measured profiles were well-described by the Gaussians and showed that the light-sheet thickness was less than 1 μm for both excitation wavelengths (FWHM_488 nm_ = 890 nm and FWHM_561 nm_ = 940 nm). Since for this measurement the light sheet plane was perpendicular to the image plane, there was some background fluorescence from fluorophores excited above and below the image plane. This out-of-focus contribution—not present when the light sheet plane coincides with the image plane—explains the deviations from a Gaussian at a larger distance from the center. These deviations could be modeled by a constant offset. The total field of view was determined by the width of the cantilever (35 μm) and length of the light-sheet. The length can be approximated to be 2× the Rayleigh length or about 10 μm. By changing the dimension of the incident laser beams with Iris 1, the thickness (FWHM) of the light sheet can be increased up to 2.5 μm resulting in an increased light-sheet length of up to 50 μm. The focus position of the light sheet, i.e. its most narrowest point, can be adjusted laterally using the electrically tunable lens ETL1. Overall, the reflected light sheet dimensions are comparable to previous ones [16] and can be varied in a certain range according to the application.

**Fig 2.**
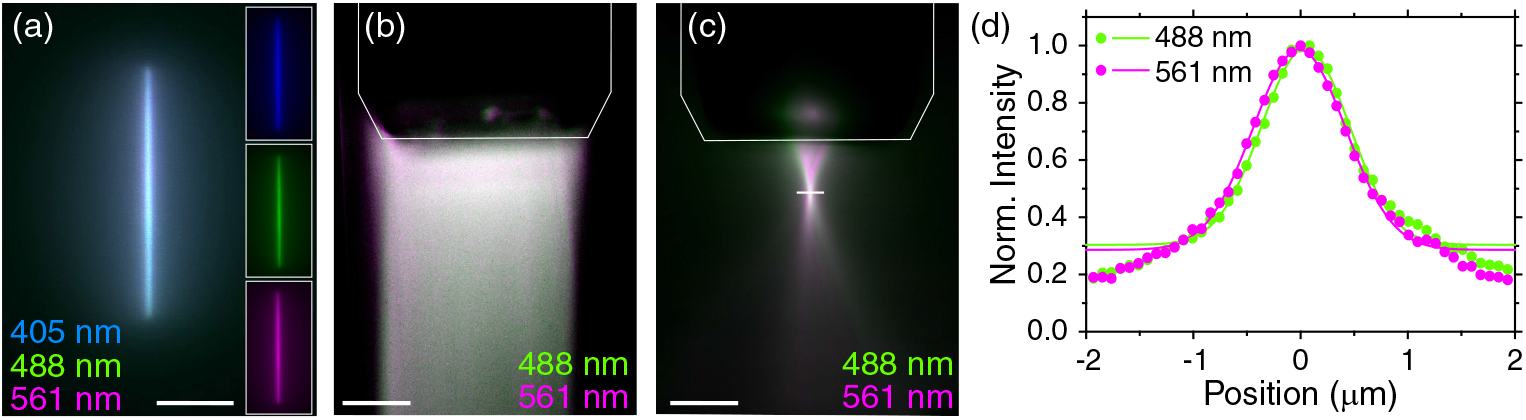
Characteristics of the light-sheet. (a) Cross-section image of the three co-aligned lasers forming the light sheet without cantilever. Insets show the individual laser cross sections. (b,c) Images of excited two-color fluorescent solution using the cantilever-reflected light-sheet. In (c) the cylindrical lens was rotated by 90° resulting in a 90° rotated light-sheet viewed from the side. The tip of the cantilever is marked by thin white lines. Scale bar: 10 μm. (d) Intensity profiles (circles) of the beam waists (white line in (c)) for the 488 nm and 561 nm laser fitted with Gaussian functions (lines, fit region: ± 1.3 μm).

### 3.2. Tracking of single molecules in GUV membranes

To demonstrate single-molecule sensitivity and the high SNR of the reflected light-sheet microscope, we measured the diffusion coefficient of single fluorescent lipids (rhodamine-PE) in a GUV (Fig. 3). Since the diffusion of these lipids was limited to the spherical membrane surface of the GUV, we positioned the light-sheet at the top of the GUV about 10 μm away from the glass surface. In this manner, the detected diffusion is approximately limited to the GUV cap and two dimensions. At a sufficiently low concentration of fluorescently labeled lipids, single lipids and their diffusive motion could be imaged (Fig. 3a, see Visualization 1). The images were acquired at a rate of 67 frames/s (15 ms per frame) allowing to track single molecules (Fig. 3b). From the single-molecule trajectories, we calculated the ensemble-averaged mean squared displacement (MSD, Fig. 3c). A least-square linear fit (4*D*τ + E) resulted in a diffusion coefficient *D* of 1.63 ± 0.03 μm^2^/s. The parameter *∈* = 4σ^2^ was 0.114 ± 0.007 μm^2^ corresponding to a localization uncertainty σ of 170 ± 40 nm [32]. Since molecules diffuse a significant distance during the integration time of a single frame, we expect the localization precision to be smaller for stationary molecules. As a control measurement, we measured the MSD of the same fluorescent lipids diffusing in a supported lipid bilayer using TIRFM (data not shown). The resulting diffusion coefficient was about 0.15 μm^2^/s. This coefficient is roughly 10× smaller compared to the one we measured in GUV caps, but is in agreement with previous work that showed that under identical conditions the lipid diffusion coefficient in GUVs is much larger than in supported lipid bilayers of identical composition [33]. Since the solid support influences the dynamics of the lipid bilayers, GUVs are a more realistic model to study the dynamics in membranes [34]. Additionally, it was shown that the diffusion coefficient of membrane proteins does not only depend on the protein size and the viscosity of the membrane and surrounding medium but also on the membrane shape and tension [35]. Therefore, it is important to provide a tool that can resolve the spatial and temporal dynamics of single membrane proteins on free-standing lipid bilayers. In addition, we determined the SNR to be about 10 (fluorophore signal divided by the background level of the camera dark offset). Our SNR was 4× higher compared to epifluorescence microscopy and only 2× lower compared to TIRFM. Thus, the high SNR of the reflected light-sheet enables single-molecule studies at micrometer distances from a glass surface.

**Fig. 3.**
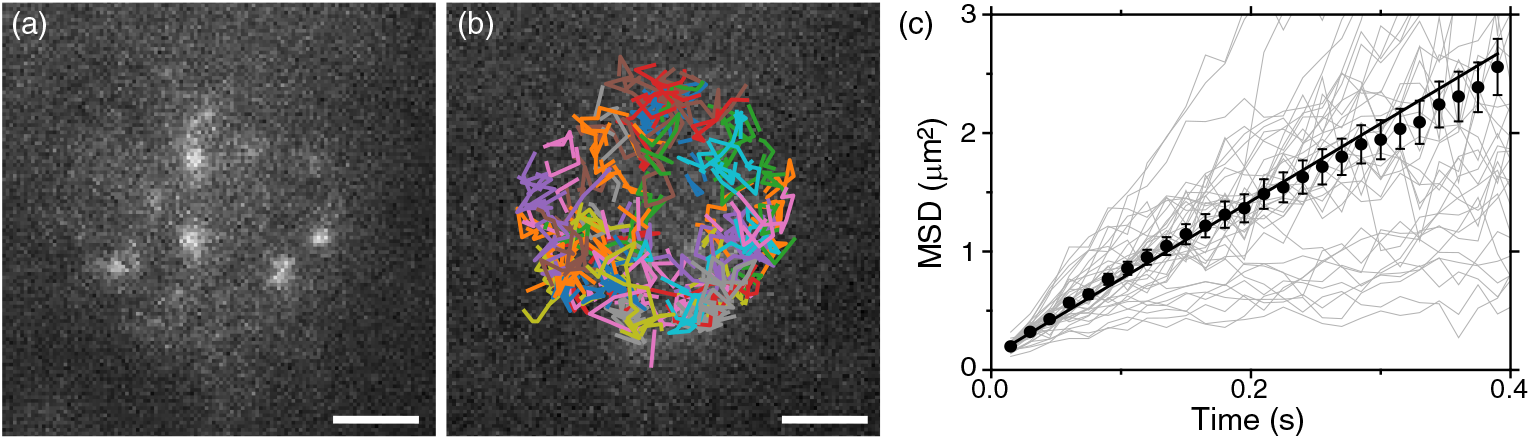
Single molecule detection of lipids diffusing in the spherical cap of a GUV. (a) Image of the GUV cap showing single lipid molecules (rhodamine-PE, see Visualization 1). Scale bar: 2 μm. (b) Single-molecule tracks. (c) Mean squared displacement (MSD) of the single traces (*N* = 39, light gray) detected in (b). Ensemble average MSD (black symbols) with standard error of the mean (SEM) and weighted linear fit.

### 3.3. Fast 3D scanning

Axial scanning of the light sheet using active optics enables fast 3D imaging without moving the sample or the objectives (Fig. 4). The imaging speed is mainly limited by the integration time of the camera. For fast scanning of the light sheet through the sample, we implemented a piezo tilt mirror in a conjugate plane to the back focal plane of the illumination objective (Fig. 4a). By tilting the piezo mirror, the laser beams will be displaced laterally in the sample. The cantilever mirror converts this lateral displacement to an axial variation of the excitation plane (*z*-axis). To match the imaging plane of the camera with the *z* position of the light sheet, we employed another electrically tunable lens (ETL2 in Fig. 1 and 4a). For fast imaging, this tunable lens needs to be synchronized with and matched in calibrated *z*-motion amplitude to the piezo tilt mirror. As a test sample with a defined geometry, we chose a spherical, fluorescently labeled GUV. To image GUV cross sections, we applied a certain set point voltage to the piezo tilt mirror illuminating only a defined plane of the GUV. Then, we adjusted the current of the tuneable lens until the cross-section was in focus. We repeated this process for different set point voltages and, in this manner, recorded a stack of cross sections at different *z* positions of the GUV (Fig. 4b). Because of the spherical geometry, we could calculate the difference in height between the individual cross sections from their respective radii. Thus, we could calibrate both the piezo tilt mirror and tuneable lens and synchronize their motion. Calibration revealed that the image *z* position depended linearly on both the voltage of the piezo tilt mirror as well as the current of the tuneable lens (Fig. 4c). This linear dependence facilitated an easy and fast automatic scanning of the sample by simultaneous adjustment of both the voltage of the piezo tilt mirror and current of the tuneable lens. The scanning speed of the piezo tilt mirrors (2 kHz), the response time of the electrically tunable lens (10 ms), and the minimal integration time of the camera (5 ms, for 2 × 2 binning) were fast enough so that the acquisition rate was limited solely by the photon budget of the sample.

**Fig. 4.**
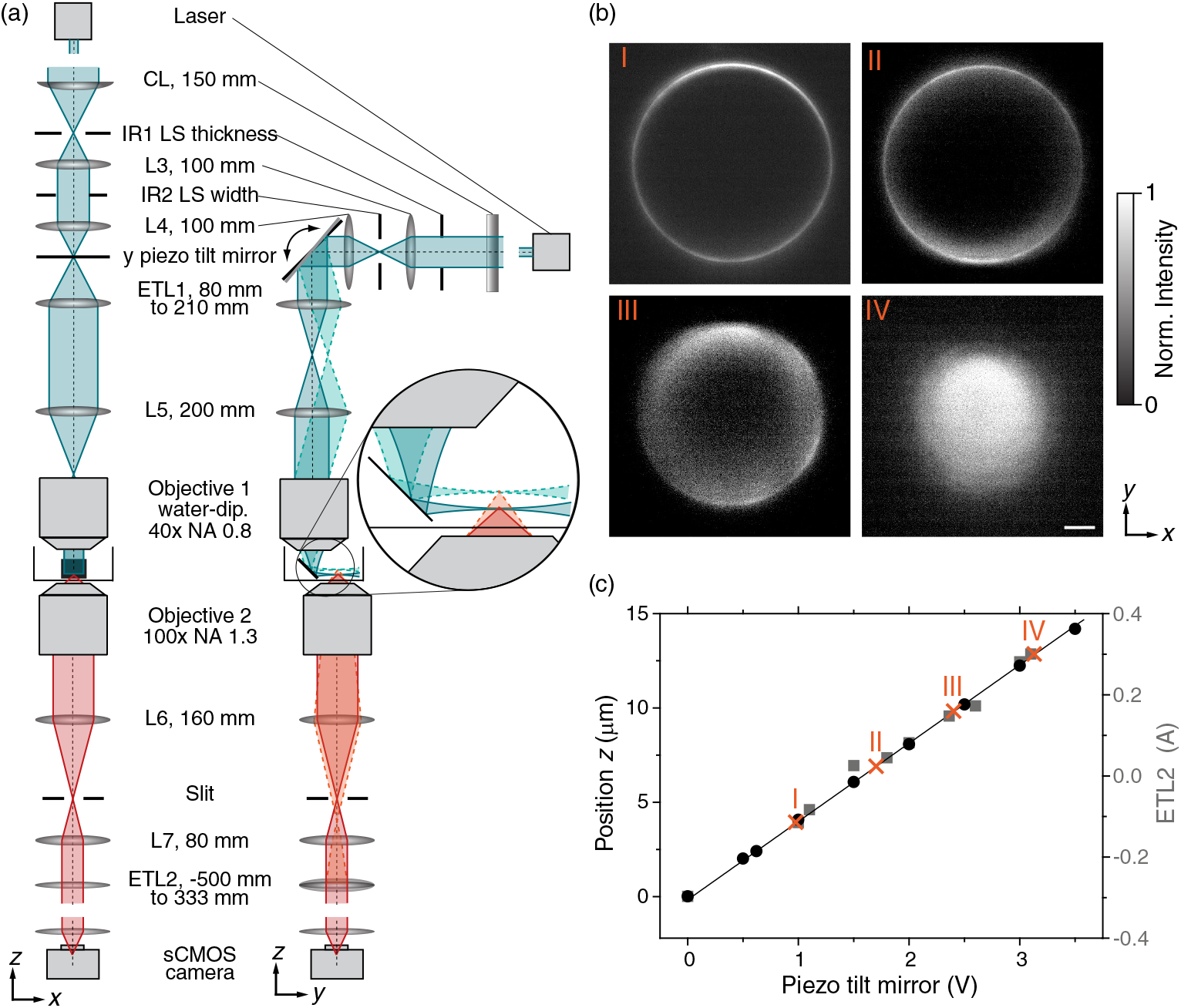
Active optics calibration for 3D imaging. (a) Schematics illustrating the two planes normal to the orientation of the optical axis, the *x*-*z* and *y*-*z* plane, respectively. A tilt of the piezo mirror results in a *y*-axis translation of the light sheet in the sample. The cantilever mirror converts this lateral *y*-motion to a vertical *z* shift of the light sheet. At the same time, an electrically tuneable lens, ETL2, is employed to adapt the image plane of the detection objective. (b) Images of a GUV, labeled with DiO, at different *z*-positions. Scans were carried out with a theoretical step size Δ*z* of 3 μm. Scale bar: 5 μm (laser output power: 100 mW, λ: 488 nm, exposure time: 25 ms). (c) Calibration of *z*-scan parameters. The *z* position of the light sheet (black symbols, left axis) depended linearly on the applied voltage on the piezo tilt mirror. Additionally, the *z* position of the detection plane depended linearly on the focal length and current of the ETL2 (gray symbols, right axis). To match the light sheet plane with the image plane, both the piezo tilt mirror and the ETL2 were moved with a constant relative factor of 0.2 (A/V). I, II, III and IV correspond to the images in (b).

To demonstrate fast, simultaneous two-color 3D imaging, we scanned through the hemisphere of a GUV labeled with both a fluorescent lipid and a membrane dye, rhodamine-PE and DiO, respectively, that were simultaneously imaged in the red and green channel (Fig. 5, see Visualization 2 and 3). The GUV diameter was ≈ 23 μm. Although the length of the light-sheet was only ≈ 10 μm, the image stack, recorded in 660 ms, could be used to reconstruct the 3D GUV structure. The reconstruction clearly shows the spherical shape of the GUV (Fig. 5a,b).

**Fig. 5.**
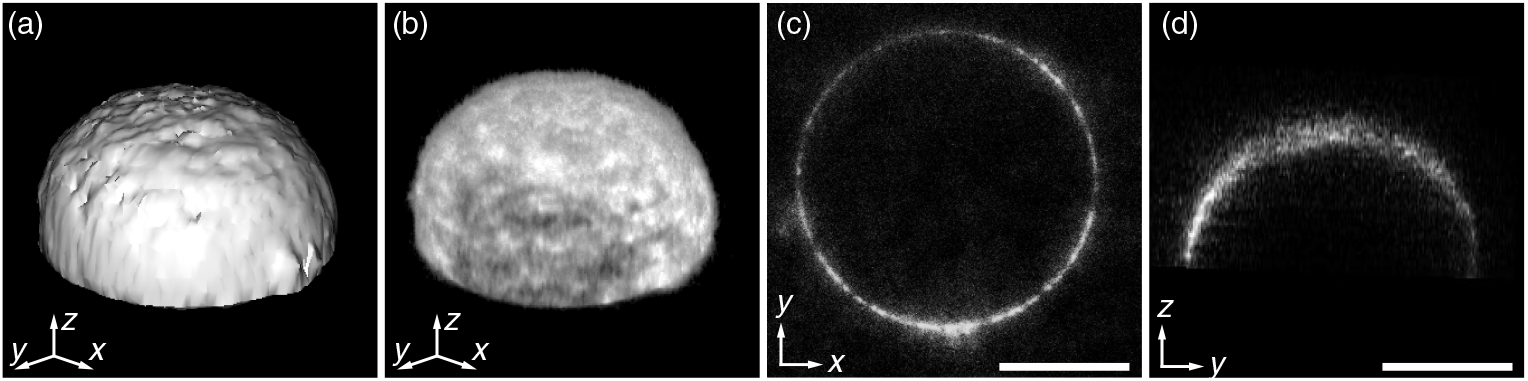
3D scan of a GUV (Ø 23 μm) labeled with DiO and rhodamine-PE. (a) Reconstructed surface plot (b) volume plot of the acquired *z*-stack. (c) *x*-*y* and (d) *x*-*z* cross-section of the GUV shown in (a,b). For reconstruction only the red-channel (rhodamine-PE) was used. Scale bar: 10 μm(laser output power: 100 mW, λ: 561 nm, scan height: 11 μm, frames: 44, step height: 0.25 μm, exposure time: 15 ms).

A cross-section through the 3D reconstructed GUV also demonstrates the high SNR and low amount of out-of-focus fluorescence (Fig. 5c,d). The scan was recorded with a step height Δ*z* of 0.25 μm, which is roughly 4× smaller than the light-sheet thickness (FWHM). In principle, 2× faster scanning could be achieved by increasing the step height to one half of the light sheet thickness in accordance with Nyquist-Shannon sampling theorem. The exposure time for each frame was 15 ms, sufficiently short to observe the diffusion of single fluorescent lipids in the cap (see Visualization 1). Note that in Fig. 5, the fluorescent lipid concentration was 10× higher compared to Fig. 3. Therefore, single molecules could not be tracked, but regions of higher and lower fluorophore density can be seen. After scanning the GUV for a second time, we did not observe any change in the fluorescent intensity, yet (data not shown), indicating that the thin light-sheet in combination with fast scanning also minimizes photobleaching. Overall, the synchronized and calibrated active optics allowed for fast, simultaneous 3D imaging of two colors with single-molecule sensitivity.

## 4. Conclusion

In this study, we presented a reflected light-sheet microscope optimized for fast 3D two-color imaging. The combination of a thin light sheet (< 1μm) and a high NA detection objective, allowed us to image and track the diffusive motion of single lipid molecules in a GUV cap. Fast 3D imaging is achieved by implementation of a piezo tilt mirror for *z*-scanning of the light sheet and an electrically tunable lens for focus adjustment. Calibration of the active optics allowed fast automatic scanning of samples. Faster scanning could be achieved by choosing a smaller region of interest on the camera for recording. The overall volume scan rate is a compromise between the number and distance between image planes, i.e. the total *z* height, camera exposure related the available laser power, the number of pixels read-out from the camera, i.e. the field of view, and the scanning speed of the active optics. Here, we used fixed increments for a new *z* position. We expect that the scanning speed can be increased by continuously varying the *z* position. Also, an optimized input signal for the electrically tunable lens may speed up the scanning process significantly [36]. Overall with the current implementation, 3D image stacks can be recorded sufficiently fast to investigate the complex dynamics inside single cells. Moreover, due to the implementation of the 405 nm laser, photoactivation experiments are feasible. Thus, this method has great potential for super-resolution imaging and 3D tracking of single molecules in living cells on a millisecond time scale.

## Supporting information

Visualization 1

Visualization 2

Visualization 3

## Supplement

**Visualization 1** Movie of fluorescent lipids (rhodamine-PE) diffusing in the cap of a GUV. The tracked lipids are shown in Fig. 3b. Movie frame rate: 3.5× slower than real time. Scale bar: 2 μm (laser output power: 100 mW, λ: 561 nm, frames: 500, exposure time: 15 ms).

**Visualization 2** One-color (red-channel, rhodamine-PE) *z* scan through a DiO and rhodamine-PE labeled GUV that was used for the 3D reconstruction shown in Fig. 5. The color code represents different *z* heights. Scale bar: 10 μm. Movie frame rate: 4.5× slower than real time (laser output power: 100 mW, scan height: 11 μm, frames: 44, step height: 0.25 μm, exposure time: 15 ms).

**Visualization 3** Two-color *z* scan through the GUV shown in Fig. 5. The GUV was labeled with two dyes (magenta: rhodamine-PE, green: DiO) and simultaneously excited with 488 nm and 561 nm. The two colors were imaged on the sCMOS camera at the same time. Movie frame rate: 4.5× slower than real time. Scale bar: 10 μm (laser output power of both lasers: 100 mW, scan height: 11 μm, frames: 44, step height: 0.25 μm, exposure time: 15 ms).

## Author contributions

E.S. and A.J. designed research; M.B., S.A.S., Q.T.D., G.H., S.D. and A.J. built and designed the microscope setup; A.J., S.A.S. and M.B. performed measurements; A.J. and E.S. wrote the manuscript.

## Acknowledgments

We thank Michael Bugiel for discussions and critical reading of the manuscript. We would like to thank Lars Lüder for first scanning experiments and Tobias Jachowski for help with Python programming.

## References and links

1. R. M. Power and J. Huisken, “A guide to light-sheet fluorescence microscopy for multiscale imaging,” Nat. Methods 14, 360–373 (2017).

2. O. E. Olarte, J. Andilla, E. J. Gualda, and P. Loza-Alvarez, “Light-sheet microscopy: a tutorial,” Adv. Opt. Photonics 10, 111 (2018).

3. J. Huisken, J. Swoger, F. D. Bene, J. Wittbrodt, and E. H. K. Stelzer, “Live embryos by selective plane illumination microscopy,” Science 305, 13–16 (2004).

4. P. J. Keller, A. D. Schmidt, J. Wittbrodt, and E. H. K. Stelzer, “Digital scanned laser light-sheet fluorescence microscopy (DSLM) of zebrafish and Drosophila embryonic development,” CSP Protocols 2011, 1235–1243 (2011).

5. Y. S. Hu, M. Zimmerley, Y. Li, R. Watters, and H. Cang, “Single-molecule super-resolution light-sheet microscopy,” ChemPhysChem 15, 577–586 (2014).

6. A.-K. Gustavsson, P. N. Petrov, and W. E. Moerner, “Light sheet approaches for improved precision in 3D localization-based super-resolution imaging in mammalian cells [Invited],” Opt. Express 26, 13122 (2018).

7. F. O. Fahrbach, P. Simon, and A. Rohrbach, “Microscopy with self-reconstructing beams,” Nat. Photonics 4, 780–785 (2010).

8. T. A. Planchon, L. Gao, D. E. Milkie, M. W. Davidson, J. A. Galbraith, C. G. Galbraith, and E. Betzig, “Rapid three-dimensional isotropic imaging of living cells using Bessel beam plane illumination,” Nat. Methods 8, 417–423 (2011).

9. A. B. Kashekodi, T. Meinert, R. Michiels, and A. Rohrbach, “Miniature scanning light-sheet illumination implemented in a conventional microscope,” Biomed. Opt. Express 9, 4263 (2018).

10. A.-K. Gustavsson, P. N. Petrov, M. Y. Lee, Y. Shechtman, and W. E. Moerner, “3D single-molecule super-resolution microscopy with a tilted light sheet,” Nat. Commun. 9, 123 (2018).

11. M. Tokunaga, N. Imamoto, and K. Sakata-Sogawa, “Highly inclined thin illumination enables clear single-molecule imaging in cells,” Nat. Methods 5, 159–161 (2008).

12. R. Galland, G. Grenci, A. Aravind, V. Viasnoff, V. Studer, and J.-B. Sibarita, “3D high- and super-resolution imaging using single-objective SPIM,” Nat. Methods 12, 641–644 (2015).

13. M. B. M. Meddens, S. Liu, P. S. Finnegan, T. L. Edwards, C. D. James, and K. A. Lidke, “Single objective light-sheet microscopy for high-speed whole-cell 3D super-resolution,” Biomed. Opt. Express 7, 2219 (2016).

14. A. Ponjavic, Y. Ye, E. Laue, S. F. Lee, and D. Klenerman, “Sensitive light-sheet microscopy in multiwell plates using an AFM cantilever,” Biomed. Opt. Express 9, 5863 (2018).

15. Y. S. Hu, Q. Zhu, K. Elkins, K. Tse, Y. Li, J. A. J. Fitzpatrick, I. M. Verma, and H. Cang, “Light-sheet Bayesian microscopy enables deep-cell super-resolution imaging of heterochromatin in live human embryonic stem cells,” Opt. Nanoscopy 2, 7 (2013).

16. J. Gebhardt, D. Suter, R. Roy, Z. Zhao, A. Chapman, S. Basu, T. Maniatis, and S. Xie, “Single-molecule imaging of transcription factor binding to DNA in live mammalian cells,” Nat. Methods 10, 421–426 (2013).

17. Z. W. Zhao, R. Roy, J. C. M. Gebhardt, D. M. Suter, A. R. Chapman, and X. S. Xie, “Spatial organization of RNA polymerase II inside a mammalian cell nucleus revealed by reflected light-sheet superresolution microscopy,” Proc. Natl. Acad. Sci. USA 111, 681–686 (2014).

18. F. Greiss, M. Deligiannaki, C. Jung, U. Gaul, and D. Braun, “Single-molecule imaging in living Drosophila embryos with reflected light-sheet microscopy,” Biophys. J. 110, 939–946 (2016).

19. F. O. Fahrbach, F. F. Voigt, B. Schmid, F. Helmchen, and J. Huisken, “Rapid 3D light-sheet microscopy with a tunable lens,” Opt. Express 21, 21010 (2013).

20. J. Jiang, D. Zhang, S. Walker, C. Gu, Y. Ke, W. H. Yung, and S.-c. Chen, “Fast 3-D temporal focusing microscopy using an electrically tunable lens,” Opt. Express 23, 24362 (2015).

21. F. Cella Zanacchi, Z. Lavagnino, M. Perrone Donnorso, A. Del Bue, L. Furia, M. Faretta, and A. Diaspro, “Live-cell 3D super-resolution imaging in thick biological samples,” Nat. Methods 8, 1047–1049 (2011).

22. G. L. Hermsdorf, S. A. Szilagyi, S. Rösch, and E. Schäffer, “High performance passive vibration isolation system for optical tables using six-degree-of-freedom viscous damping combined with steel springs,” Rev. Sci. Instrum. p. 015113 (2019).

23. S. Simmert, M. K. Abdosamadi, G. L. Hermsdorf, and E. Schäffer, “LED-based interference-reflection microscopy combined with optical tweezers for quantitative three-dimensional microtubule imaging,” Opt. Express 26, 14499 (2018).

24. M. Mahamdeh, S. Simmert, A. Luchniak, E. Schäffer, and J. Howard, “Label-free high-speed wide-field imaging of single microtubules using interference reflection microscopy,” J. Microsc. 272, 60–66 (2018).

25. A. Edelstein, N. Amodaj, K. Hoover, R. Vale, and N. Stuurman, “Computer control of microscopes using manager,” in “Current Protocols in Molecular Biology,” (John Wiley & Sons, Inc., Hoboken, NJ, USA, 2010), SUPPL. 92, pp. 1–17.

26. M. I. Angelova, S. Soléau, P. Méléard, F. Faucon, and P. Bothorel, “Preparation of giant vesicles by external AC electric fields. Kinetics and applications,” Progr. Colloid Polym. Sci. 89, 127–131 (1992).

27. A. J. García-Sáez, J. Ries, M. Orzáez, E. Pérez-Payà, and P. Schwille, “Membrane promotes tBID interaction with BCL(XL),” Nat. Struct. Mol. Biol. 16, 1178–1185 (2009).

28. J. D. Unsay, K. Cosentino, and A. J. García-Sáez, “Atomic force microscopy imaging and force spectroscopy of supported lipid bilayers,” J. Vis. Exp. p. e52867 (2015).

29. S. Sudhakar, T. J. Jachowski, M. Kittelberger, A. Maqbool, G. L. Hermsdorf, M. K. Abdosamadi, and E. Schäffer, “Supported solid lipid bilayers as a platform for single-molecule force measurements,” Nano Lett. 19, 8877–8886 (2019).

30. J. Schindelin, I. Arganda-Carreras, E. Frise, V. Kaynig, M. Longair, T. Pietzsch, S. Preibisch, C. Rueden, S. Saalfeld, B. Schmid, J.-Y. Tinevez, D. J. White, V. Hartenstein, K. Eliceiri, P. Tomancak, and A. Cardona, “Fiji: an open-source platform for biological-image analysis,” Nat. Methods 9, 676–682 (2012).

31. D. B. Allan, T. Caswell, N. C. Keim, and C. M. van der Wel, “trackpy: Trackpy v0.4.1,” (2018).

32. X. Michalet, “Mean square displacement analysis of single-particle trajectories with localization error: brownian motion in isotropic medium,” Phys. Rev. E 82, 041914 (2010).

33. M. Przybylo, J. Sýkora, J. Humpolíčková, A. Benda, A. Zan, and M. Hof, “Lipid diffusion in giant unilamellar vesicles is more than 2 times faster than in supported phospholipid bilayers under identical conditions,” Langmuir 22, 9096–9099 (2006).

34. R. Macháň and M. Hof, “Lipid diffusion in planar membranes investigated by fluorescence correlation spectroscopy,” Biochim. Biophys. Acta, Biomembr. 1798, 1377–1391 (2010).

35. F. Quemeneur, J. K. Sigurdsson, M. Renner, P. J. Atzberger, P. Bassereau, and D. Lacoste, “Shape matters in protein mobility within membranes,” Proc. Natl. Acad. Sci. USA 111, 5083–5087 (2014).

36. D. Iwai, H. Izawa, K. Kashima, T. Ueda, and K. Sato, “Speeded-up focus control of electrically tunable lens by sparse optimization,” Sci. Rep. 9, 12365 (2019).

